# kimma: flexible linear mixed effects modeling with kinship covariance for RNA-seq data

**DOI:** 10.1101/2022.10.10.508946

**Authors:** Kimberly A Dill-McFarland, Kiana Mitchell, Sashank Batchu, R Max Segnitz, Basilin Benson, Tomasz Janczyk, Madison S Cox, Harriet Mayanja-Kizza, W Henry Boom, Penelope Benchek, Catherine M. Stein, Thomas R Hawn, Matthew C Altman

## Abstract

We introduce kimma (Kinship In Mixed Model Analysis), an open-source R package for flexible linear mixed effects modeling of RNA-seq including covariates, weights, random effects, covariance matrices, and fit metrics. In simulated datasets, kimma detects differentially expressed genes (DEGs) with similar specificity, sensitivity, and computational time as limma unpaired and dream paired models. Unlike other software, kimma supports covariance matrices as well as fit metrics like AIC. Utilizing genetic kinship covariance, kimma revealed that kinship impacts model fit and DEG detection in a related cohort. Thus, kimma equals or outcompetes current DEG pipelines in sensitivity, computational time, and model complexity.

## Background

Transcriptomics provide a powerful method to capture genome-wide RNA expression at relatively low costs. Methods such as microarray and RNA-seq allow for in-depth assessment across populations and time scales relevant to research in human health, disease, ecology, and others. Transcriptomics studies tend to have experimental designs with many more measures (e.g. genes) than observations (e.g. subjects or libraries). Thus, subsequent analyses require multiple comparison correction, which can obscure true biological results in an effort to reduce Type I error [1]. In addition, numerous or complex covariates can be necessary to correct for diverse population features and designs can include repeated measures, paired sampling, or blocking. Together, these features require diverse statistical methods and robust model fit assessments not fully supported by current transcriptomic computational tools.

Presently, there are a variety of analysis pipelines for the identification of differentially expressed genes (DEGs) in bulk transcriptomic datasets [2]. DEGs represent one of the major results from RNA-seq experiments and drive discoveries in this space. Among these pipelines, limma [3] and DESeq2 [4] are among the most popular, each with well over 10,000 citations since their publications in 2015 and 2014, respectively. Limma utilizes simple linear regression and empirical Bayes statistics to assess differential gene expression. This R package also supports gene-specific weights to account for library quality [5] and a pseudo-random effect to account for paired sample designs or repeated measures. Others have expanded on the limma framework with dream in the VariancePartition R package to incorporate true random effects to more accurately account for paired designs as well as support multiple blocking variables in complex mixed effects designs [6]. In contrast, DESeq2 employs negative binomial regression to assess differential expression. This statistical framework does not require gene-level weights and can fit small datasets with large ranges or outliers. DESeq2, however, does not support random effects and cannot account for paired designs.

While current RNA-seq analysis tools can accommodate a wide range of univariate and multivariate predictors as well as simple random effects, none support the inclusion of complex random effects like covariance matrices, which consist of more than one measure per observation. Thus, these effects cannot be accounted for using current RNA-seq tools. One potential covariance of interest is kinship, a summative measure of genetic relatedness between pairs of individuals. Kinship more accurately describes genetic relatedness than social constructs like race and ethnicity [7], which would be used as fixed effects in modeling, and can explain significant variation in RNA-seq data, particularly in related cohorts [8].

Here, we present kimma for kinship in mixed model analysis. Kimma is an open-source R package that provides flexible linear mixed effects modeling for bulk RNA-seq data including univariate, multivariate, random, and covariance random effects as well as gene-level weights. Building on well-tested statistical packages in R such as stats [9] and lme4 [10], kimma utilizes a single function, kmFit, for modeling, ensuring consistent syntax, inputs, and outputs. Moreover, kimma provides post-hoc pairwise tests, model fit metrics like AIC, and fit warnings on a per gene basis. Kimma’s outputs can also be integrated into our sister R packages in the BIGverse for visualization of results as well as gene set analyses.

## Results

### Overview of kimma

The kimma R package provides flexible linear modeling of RNA-seq data with a single function, kmFit. This package is open-source and freely available on GitHub (https://github.com/BIGslu/kimma). Kimma fits linear and/or linear mixed effects models incorporating gene-level weights, covariates, and/or random effects. In addition, kimma provides easy comparison of model fits with metrics such as AIC, BIC, and R-squared as well as downstream analyses including summary tables and expression quantitative loci (eQTL). The kimma outputs feed directly into our related packages such as plotting in BIGpicture and gene set analysis in SEARchways. Collectively, these R packages work together in the meta-package BIGverse (Figure 1).

**Figure 1.**
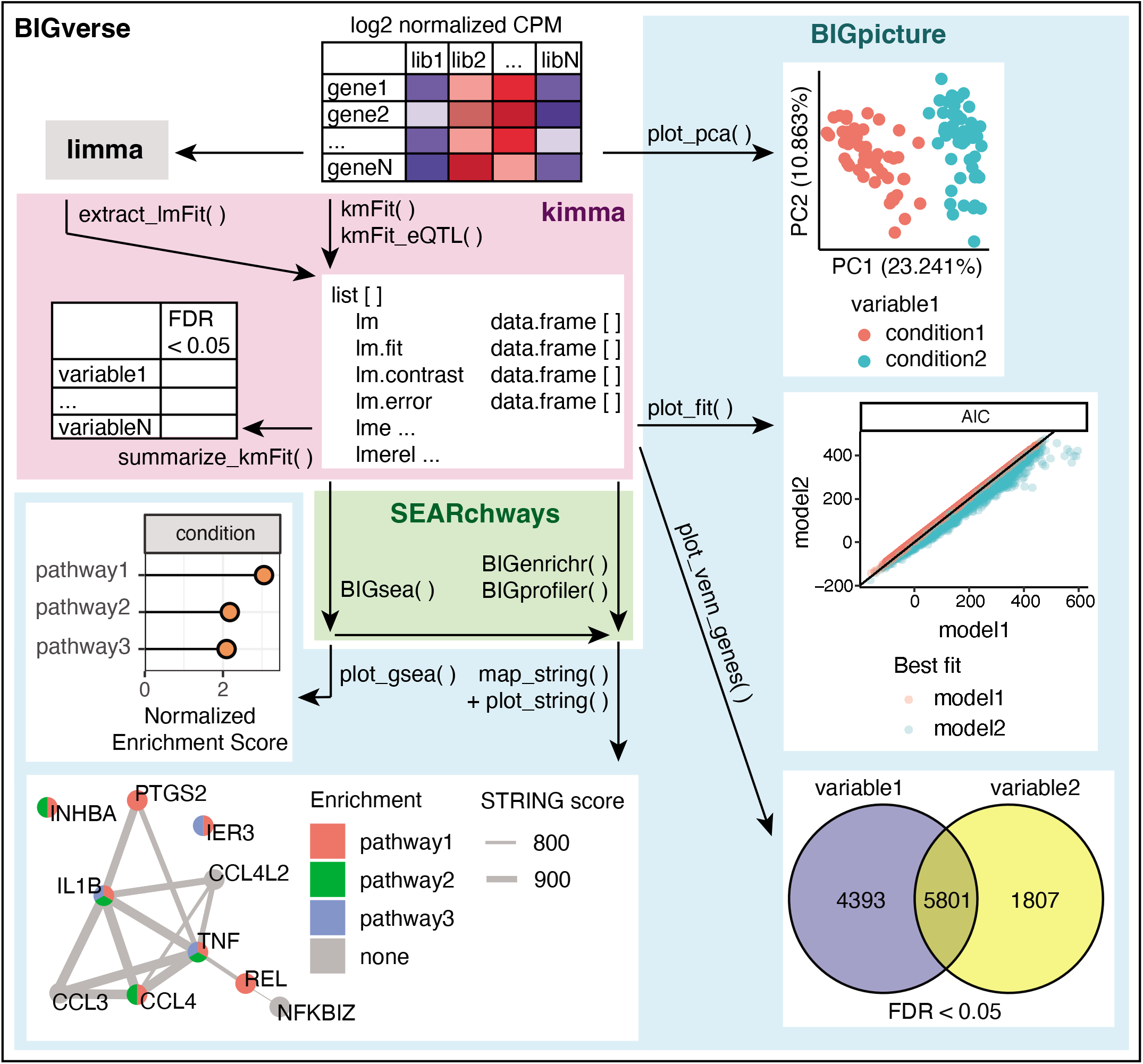
BIGverse workflow. Log2 normalized counts per million (CPM) gene expression are input into the BIGverse pipeline. Prior to modeling, variables of interest can be explored by PCA using BIGpicture. Gene expression is then modeled in kimma with kmFit using simple linear (lm), linear mixed effects (lme), and/or lme with kinship relatedness (lmerel). Each model results in up to four data frames: estimates and significance, goodness of fit metrics, pairwise contrasts, and errors for genes that failed fitting. Model fit is assessed by metrics, such as AIC, which can be plotted in BIGpicture. Significant genes are summarized in kimma and plotted using Venn diagrams in BIGpicture. SEARchways is used for pathway analyses such as GSEA with model estimates of fold change and hypergeometric pathway enrichment of significant genes. Significant pathways are visualized as well as annotated to specific genes in STRING networks using BIGpicture. In addition, kimma provides functionality to import limma results for use in parallel analyses and visualization.

### kimma performs as well as current methods for differentially expressed gene (DEG) analysis without kinship in a simulated data analysis

To test kimma’s performance, we constructed simulated test datasets from previously published RNA-seq data which included two groups (resister [RSTR)] and latent TB infection [LTBI]) and two conditions (media and *M. tuberculosis* [Mtb] infection) in a household-based cohort of individuals with varying degrees of relatedness [8,11]. Modeling was performed in kimma, limma, dream, and DESeq2 using 100 simulated datasets of 250 DEGs with fold changes from 1 to 100% and 750 non-DEGs (Figure S1A). In unpaired model designs, kimma and limma resulted in similar sensitivity and specificity (mean AUC 0.93, Figure 2). DESeq2’s negative binomial model performed significantly worse than all unpaired kimma or limma models (mean AUC 0.57, FDR < 0.05). In paired designs, kimma’s random effect and dream’s repeated measures (mean AUC 0.98) slightly outperformed limma’s duplicateCorrelation pseudo random effect (mean AUC 0.95) and greatly outperformed DESeq2’s fixed effect for donor (mean AUC 0.47, FDR < 0.05). Within a model type and software, gene-level weights had no effect on DEG detection.

**Figure 2.**
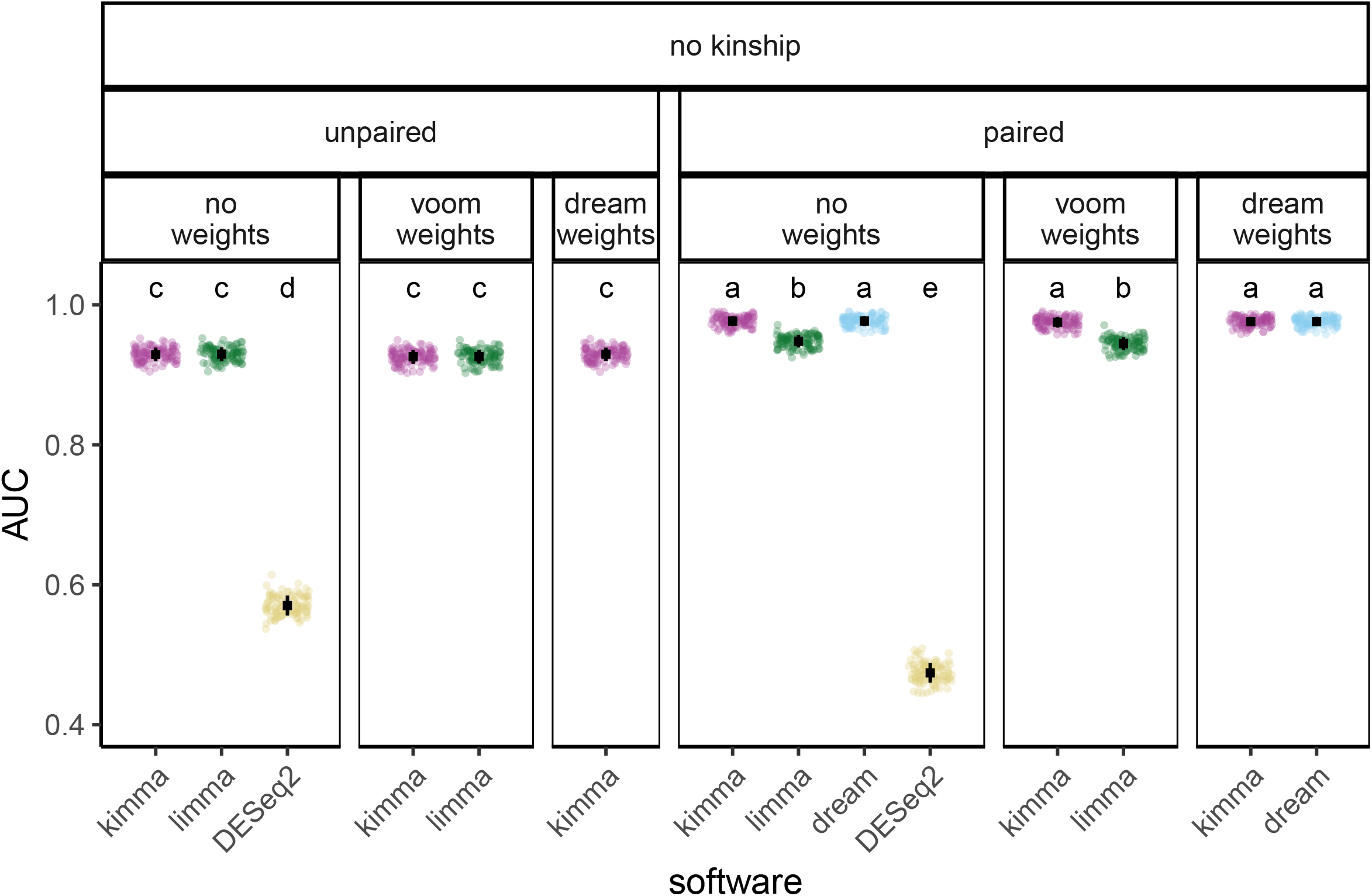
Differentially expressed genes detected across linear models and software. Area under the curve (AUC) was calculated from curves of 1-specificity to sensitivity for P-value cutoffs from 0 to 1 in 100 simulated datasets. When available, models were corrected for gene-level weights (kimma, limma, dream) and/or paired design (kimma, limma, dream, DESeq2) as indicated by panel headers. Black squares indicate means with standard deviation bars. Across all panels, letters indicate statistically different groups, TukeyHSD FDR < 0.05, in order from highest AUC (a) to lowest (e). Points are colored by software package.

### kimma’s multi-processor functions achieve runtimes similar to limma

Multiple processor functionalities in kimma, dream, and DESeq2 were compared across simulated datasets (Figure S1A) on six processors as this is a reasonable load for standard 8-core laptops. In paired model designs, kimma runtime linearly decreased from 72 seconds on 1 processor to 12 seconds on 6 processors (Figure 3). Kimma’s 6-processor speed was as fast or slightly faster than limma’s 1 processor speed for paired designs (FDR < 0.05). In contrast, dream was slower and did not reduce time linearly, taking 55 seconds on 1 processor and 26 seconds on 6 processors. DESeq2 was the slowest software for paired designs, though this method uses donor as a fixed effect, thus increasing model complexity compared to other methods. In contrast, limma outperformed all other software for speed in unpaired designs (Figure S2).

**Figure 3.**
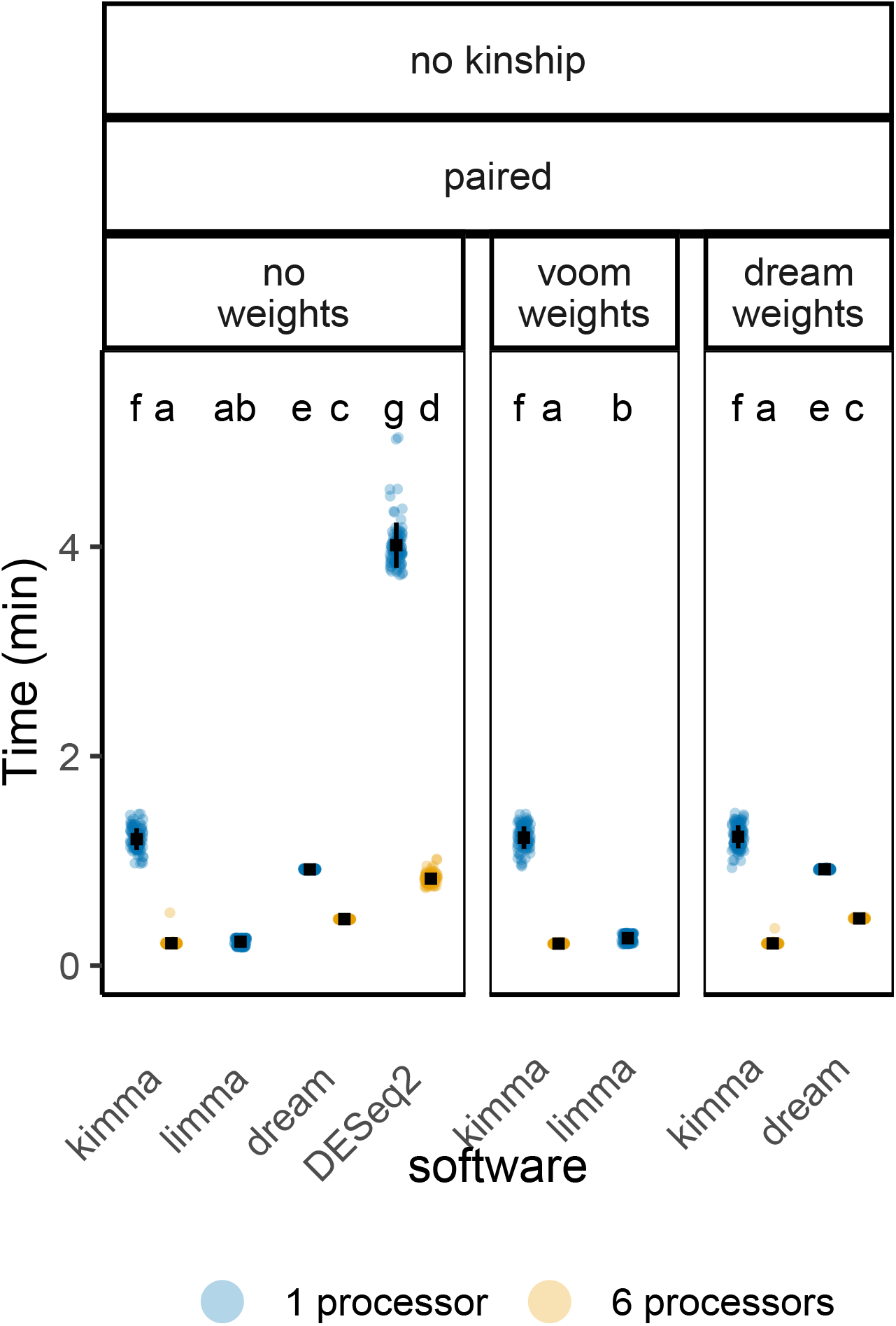
Runtime to analyze simulated RNA-seq data across paired models and software. Models were run for 100 simulated datasets of 92 samples and 1000 genes. When available, models were corrected for gene-level weights (kimma, limma, dream) and/or paired design (kimma, limma, dream, DESeq2) as indicated by panel headers. Multi-processor usage is shown for kimma, dream, and DESeq2 at 1 (blue) or 6 processors (orange). Limma does not have this option. Black squares indicate means with standard deviation bars. Across all panels, letters indicate statistically different groups, TukeyHSD FDR < 0.05, in order from shortest time (a) to longest (g).

### kimma provides additional functionalities at no cost to computational time

We next used a full-size data analysis to test whether kimma can incorporate multiple model types and other features while keeping computational time low. In a dataset of 13,972 protein-coding genes and 98 samples, model fitting and significance estimation takes less than 90 seconds (Figure S3). The addition of fit metric calculation, gene-level weights, covariates, or interaction terms has no effect on total processing time. Adding pairwise contrasts between multi-level variables or interaction terms, paired sample design as a random effect, or kinship as a covariance random effect increases computational time to between 3 and 4 minutes for this dataset. Combining kimma’s features increases processing time almost linearly as a model with all seven features increased time by a factor of six and running three models at once takes roughly the sum of time for the three models run separately.

### kimma detects more DEGs in real-world paired model designs

We next compared DEG detection in real-world data. In a dataset of Mtb-infected vs media controls in LTBI and RSTR donors [8], DEGs were modeled using the best performing methods based on simulated data analyses (unpaired group c, paired group a, Figure 2). Thus, kimma and dream were compared for paired designs (Figure 2, group a) while kimma and limma were compared for unpaired designs (Figure 2, group c).

For paired Mtb-infected and media samples (Figure S1B), the majority of DEGs were detected by all models, and gene-level weights resulted in the largest addition of 209 DEGs regardless of software (FDR < 0.05, Figure 4A). Overall, kimma with weights resulted in the most DEGs with 94 more DEGs than dream with weights, of which 50 were specific to only the kimma with weights method. At an FDR of 0.05, simulated data analysis showed kimma to have minimally but consistently higher sensitivity (max delta sensitivity 0.02) when compared to dream (Table S1). This supports the detection of more true-positive DEGs by kimma in the Mtb analysis compared to dream. In contrast, an unpaired analysis of LTBI vs RSTR (Figure S1C) resulted in 6 DEGs identified by all models and software (FDR < 0.05, Figure 4C). This is further supported by the unpaired simulated analysis where differences in sensitivity were very low and did not always support the same model (max delta sensitivity 0.008, Table S1).

**Figure 4.**
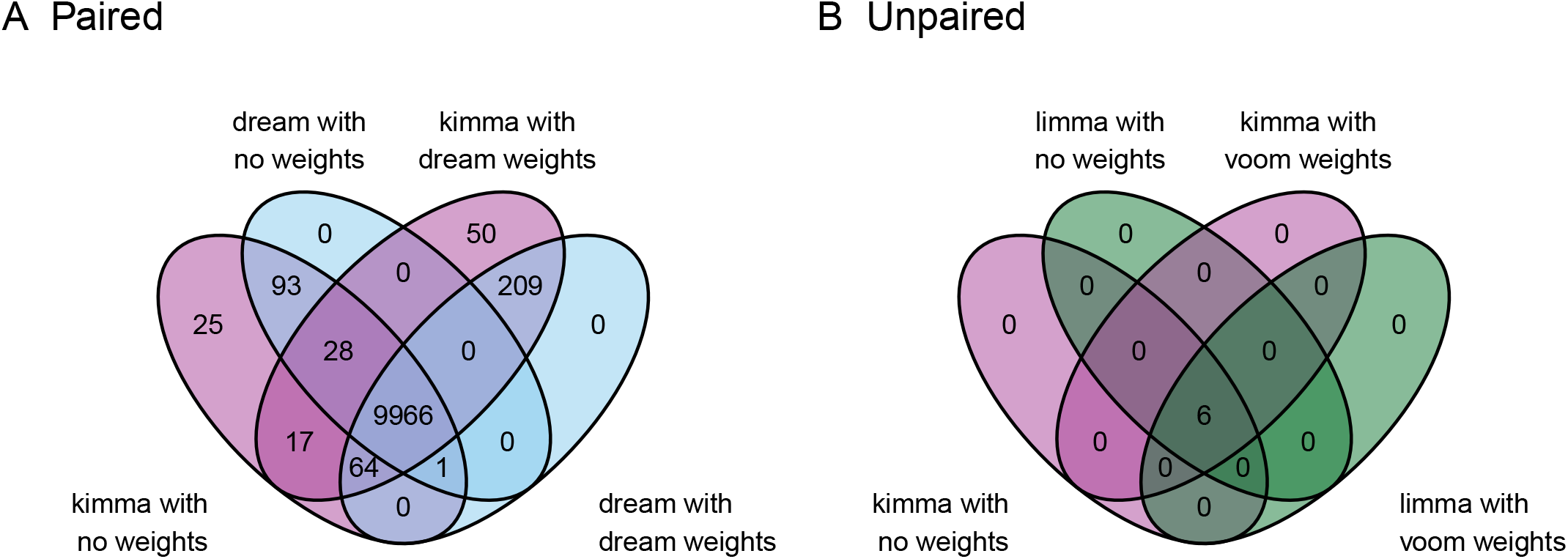
Real-world RNA-seq data analysis across linear models and software. Using the best performing models from simulated data, real-world data was assessed for the impacts of (A) Mtb infection in paired designs (Figure 2, group a) and (B) RSTR status in unpaired designs (Figure 2, group c). Venn diagrams show total differentially expressed genes (DEGs).

### Kinship improves model fit and DEG detection in paired, genetically related datasets

Unlike other RNA-seq analysis packages, kimma allows incorporation of a covariance random effect such as pairwise genetic kinship. Here, the Mtb vs media and RSTR vs LTBI data were split into unrelated (kinship < 0.125) and related subsets (at least one kinship > 0.125) (Figure S1B,C) to assess the impacts of kinship in linear mixed effects models. In the paired Mtb-infected vs media model, the addition of kinship among unrelated individuals did not impact model fit (Figure 5A,B) with only 2 genes reaching an absolute delta AIC greater than 2 (max delta AIC = 2.4). This corresponded with minimal changes in DEG detection as both models identified 99.9% of the same genes at FDR < 0.05 (Figure 5C).

**Figure 5.**
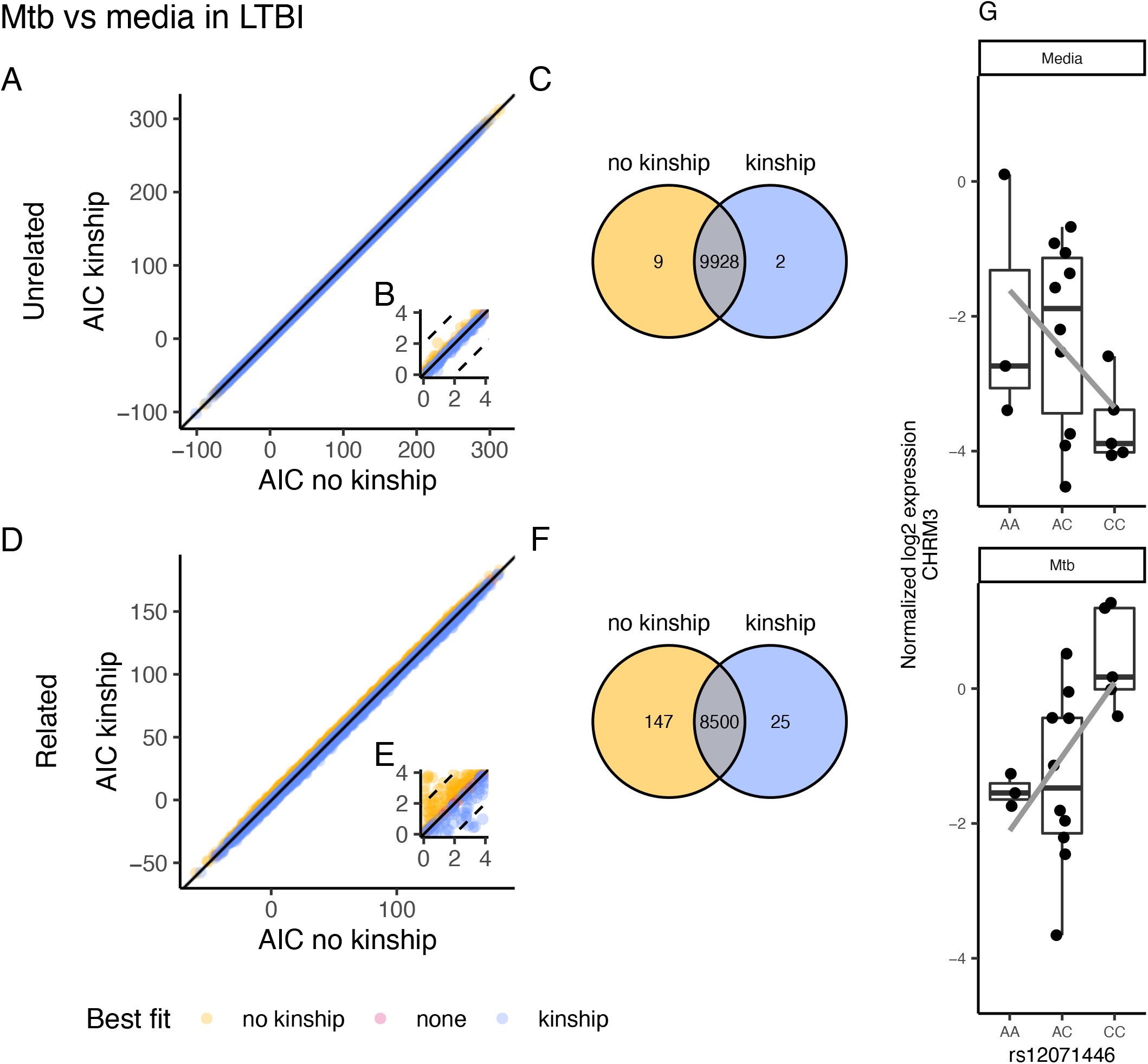
Paired design model fit and DEGs in genetically unrelated and related datasets. LTBI individuals were subset into unrelated and related subsets at a kinship cutoff of 0.125. Subsets were modeled for Mtb-infected vs media expression. (A,B) In unrelated individuals, model fit was assessed by AIC for a model with and without a kinship random effect. The solid black line indicates a 1:1 fit (e.g. no difference in AIC) and dashed lines indicate ± 2 AIC (e.g. moderate difference in AIC). The inset shows AIC from 0 to 4 highlighting genes better fit without kinship in the model (yellow) and those better fit with kinship in the model (blue). (C) DEGs were defined at FDR < 0.05. (D,E,F) Similarly, model fit and DEGs were assessed for the related subset. (G) Expression of the top Mtb:SNP cis eQTL from kinship DEGs in the related subset.

In contrast, Mtb model fit of related individuals was impacted by kinship (Figure 5 D,E) with 1247 genes better fit by > 2 AIC with kinship (max delta AIC = 6.6) and 518 better fit without kinship (max delta AIC = 6.8). This resulted in changes in DEG detection with 25 new DEGs detected in the kinship model (Figure 5F). Importantly, 91 of the 147 DEGs only detected without kinship were better fit by the model with kinship according to AIC. Thus, the addition of kinship appears to reduce Type I error by improving fit for a subset of genes.

In the unpaired RSTR vs LTBI design, kinship did not improve model fit with the majority of genes (98%) better fit without kinship in the model for both unrelated and related subsets (max delta AIC = 15, Figure S4A,C). While more DEGs were detected with kinship in the unpaired, related subset, the majority of these genes (161 of 165) were better fit by AIC without kinship in the model. Thus, the inclusion of kinship in this case may result in higher Type I error due to a poorer fitting unpaired model.

### kimma extends to other applications such as eQTL analysis

Downstream analysis of RNA-seq data often includes expression quantitative trait loci (eQTL), where single nucleotide polymorphisms (SNPs) are associated with gene expression. Kimma’s kmFit_eQTL function provides the means to test eQTLs with the same flexible modeling and fit assessment features as kmFit. As a test case, we utilized genome-wide SNP data from the same cohort to determine eQTL for DEGs identified in the related kinship model. Among the kinship model DEGs (N = 8525, Figure 5F), we identified 10020 potential within-gene cis eQTL in 3382 unique genes. In total, 289 eQTL in 253 unique genes were significant for the interaction of SNP and Mtb infection (FDR < 0.05). The top hit was genotype rs12071446 which was associated with Mtb-induced expression of the CHRM3 gene (FDR = 2.2E-7, Figure 5G). This analysis framework can be readily applied to any gene(s) of interest to determine genetic regulation of gene expression using the same robust models as DEG detection.

## Discussion

Transcriptomics is utilized in many fields to address diverse scientific questions. Identification of differentially expressed genes (DEGs) remains a cornerstone of analysis of these data, and current bioinformatic tools support a range of study designs. Here, we introduce kimma, an open-source R package that expands supported designs to include covariance random effects as well as provides metrics for robust model fit assessment. In addition, kimma provides model fit assessment and integrates with several downstream analysis and visualization functions for a more streamlined RNA-seq analysis pipeline.

Kimma does not present new statistical methods but instead, builds on well-tested linear modeling methods in R. Simple linear models use base R’s lm [9] and linear mixed effects models use lme4’s lmer [10]. Both are widely used and have been available in R for nearly 20 years. Models with covariance matrices are a new addition (2018) and utilize lme4qtl’s relmatLmer [12], which builds on the foundation of lme4. Kimma brings these methods together, codifying similar parameters such as gene-level weights, and results in consistent, organized outputs which flow into the BIGverse RNA-seq analysis pipeline including data visualization in BIGpicture, gene set analyses in SEARchways, and eQTL analysis in kimma.

In simulated paired study designs, kimma performs as well as or better than limma [3,5], dream [6], and DESeq2 [4]. Kimma’s true mixed effects modeling results in comparable sensitivity and specificity to dream, both of which outperform limma’s pseudo random effect with duplicateCorrelation. Kimma outperforms dream with shorter computational time on multiple processors. In fact, kimma’s multi-processor functionality affords runtimes on the order of minutes for full-size RNA-seq data (∼14,000 genes, 100 samples) even when multiple additional features or models are requested. These results translate to real-world paired analyses where kimma detects more DEGs than comparable models in dream. Our results suggest these are true positives as kimma also resulted in slightly higher sensitivity in simulated DEG detection.

In predefined unpaired study designs, limma remained the preferred software because while limma and kimma had similar simulated DEG detection, limma achieved faster runtimes. However, limma does not support model fit assessment through metrics such as AIC and requires multiple models to run pairwise contrasts. In addition, kimma out-performed DESeq2 with significantly better DEG detection. Thus, kimma equivalent or better DEG detection along with additional functionalities in unpaired analyses where model selection is on-going or multi-level variables such as interaction terms are of interest.

Kimma also allows for the use of covariance matrix random effects in linear modeling; these random effects are not supported by any other current method. Here, we use pairwise genetic kinship as an example, but any covariance matrix may be used. In paired study designs, kinship improved model fit and reduced Type I error in DEG detection in genetically related individuals. This effect was apparent even though the current dataset contains only sparse relatedness with 1.4% of pairs achieving third degree relatedness or higher. Model fit was not significantly impacted by kinship in unrelated individuals. In addition, kinship did not improve model fit and appeared to increase Type I error in unpaired designs. Thus, paired designs benefit from kinship inclusion and are not negatively impacted when kinship is nonsignificant. Together, these analyses highlight the advantages of kimma’s model fit assessment in RNA-seq analyses as other workflows may have readily accepted these DEGs even if the model was not best fit for the data. Poor model fit can lead to non-reproducible results and wasted resources on downstream experiments pursuing what were originally false positives.

Kimma’s flexible model design can be extended for targeted eQTL analysis. Using a simple model of genotype:Mtb infection corrected for kinship, we identified 289 eQTL among Mtb DEG in related individuals. The purpose of this analysis was not to identify novel Mtb eQTL as a full genetic analysis likely requires further covariate assessment such as population principal components, sex, etc. Instead, this showcases kimma’s ability to model more than just RNA-seq differential expression. Given kimma’s computational time, tools such as GENESIS [13] or GMMAT [14] are preferred for genome-wide eQTL analyses. However, kimma provides robust targeted analysis with the same underlying model as differential gene expression as well as yields fit metrics useful in determining covariate usage on a subset of genes prior to genome-wide GENESIS or GMMAT analyses.

Due to the limited availability of open-source RNA-seq data from paired designs with accompanying covariance matrix random effects like kinship, kimma was applied to only one real-world dataset in these analyses. While simulated datasets provided measurable sensitivity and specificity, other real-world data should be explored to better elucidate the impacts of genetic kinship or other covariances on DEG detection. We expect the use of kinship with transcriptomic data to increase, in particular as methods to extract genotype information directly from RNA-seq data become more routinely used, thereby not requiring additional biological material or monetary cost [15]. In addition, while kimma was applied to RNA-seq data in these examples, the package itself is not RNA-seq specific. Any dataset with multiple measures can be coerced into a similar format and modeled using the appropriate parameters. The user need only ensure that the data meet the requirements of the linear model being applied.

## Conclusion

Overall, kimma provides a user-friendly, single function workflow for linear model analysis of RNA-seq data. It further provides model fit and downstream analyses relevant to DEGs identified by these linear models. Users can explore kimma and its sister packages for results visualization and downstream gene set analyses in the BIGverse through open-source vignettes available at https://bigslu.github.io/kimma_vignette/

## Methods

### Tuberculosis dataset

Study recruitment [16] and RNA-sequencing experiments [8,11] were previously described. Briefly, household contacts of individuals with active pulmonary tuberculosis were followed for 8 to 10 years and classified as persistently TST-/IGRA-resisters (RSTR) or latent tuberculosis infection (LTBI). Monocytes were extracted from whole blood and cultured without stimulation (media) or with *Mycobacterium tuberculosis* strain H37Rv (Mtb) (MOI 1.0, 6 hours). Bulk RNA-sequencing was performed using Illumina 50 bp paired-end sequencing, and sequences were aligned to GRCh38 with STAR [17]. Low abundance genes were removed if they did not reach at least 1 count per million (CPM) in 5% of samples in the original dataset. Genetic kinship was determined using genotypes from the Illumina MEGAEX array (2 million SNP) with the robust KING method for identity by descent and a genetic relationship matrix (GRM) [18].

### Simulated data generation

In order to maintain the paired sample design, 100 simulated DEGs datasets were created from the uninfected, media samples from the RSTR/LTBI tuberculosis dataset. In total, 1000 random genes were selected and copied for simulated samples. Then, 50 DEGs were created at each fold change of 1, 5, 10, 50, and 100%, and random error of ± 5% was introduced to all genes.

### limma and dream models

Gene expression was modeled as trimmed mean of means (TMM) normalized log2 CPM. For limma (v3.50.3 [3]), linear models were fit, and p-values were estimated using the empirical Bayes method. Pseudo-paired sample design was estimated using mean correlation of expression between paired samples across all genes (duplicateCorrelation). For dream (variancePartition v1.24.0 [6]), only paired sample designs were run as its unpaired designs are equivalent to limma. Linear mixed effects models were fit blocked by donor, and p-values were estimated using the empirical Bayes method. Where specified, gene-level quality weights were calculated using voomWithQualityWeights for limma or dreamWithQualityWeights for dream and incorporated in the models.

### DESeq2 models

Gene expression was modeled as unnormalized counts as recommended by the software. For DESeq2 (1.34.0 [4]), negative binomial general linear models including total library size and dispersion estimated were fit, and p-values were estimated using the Wald test. Pseudo-paired sample design was estimated by including donor as the first main effect term. DESeq2 does not support gene-level quality weights.

### kimma models

Gene expression was modeled as TMM normalized log2 CPM. For kimma (v1.2.0), linear models were fit, and p-values were estimated using restricted maximum likelihood (REML). Simple linear regression was performed with base R stats::lm, mixed effects for paired sample designs with lme4::lmer, and mixed effects for paired sample designs with kinship with lme4qtl::relmatLmer. As with limma, gene-level quality weights calculated using voomWithQualityWeights were incorporated for some models. Though not applicable in these analyses, kimma can also fit pairwise contrasts within main model terms using estimated marginal means in the emmeans package [19].

### Simulated data analysis

Simulated differentially expressed genes were modeled against their corresponding media samples. Within each of 100 simulated datasets, sensitivity was defined as true positive / (true positive + false negative) and specificity as true negative / (true negative + false positive). Area under the curve (AUC) was calculated for 1 -specificity by sensitivity at p-value cutoffs in increments of 0.001 from 0 to 0.01 plus increments of 0.01 from 0.01 to 1. Time trials were run on Amazon Web Services (AWS, 48 vCPU, 192 GB RAM) with 1 or 6 processors as appropriate and nothing else running.

### Tuberculosis analysis

In total, 13,972 pass-filter genes (see [8]) were modeled for media vs Mtb-infected in paired LTBI samples (N = 92) as well as RSTR vs LTBI in unpaired Mtb-infected samples (N = 98). Analyses were completed on the entire dataset as well as subsets of unrelated (kinship < 0.125, N = ^55, 62^) and related samples (at least one kinship > 0.125, N = ^37, 36^). Genes found to be significant for Mtb vs media in the related subset with kinship correction were extracted from genetic data with SNPs found on both the MEGAEX and Omni chip [18]. In total, 3382 of these 8525 DEGs had at least 1 pass-filter SNP within the gene transcript region (cis) to yield 10020 unique SNP-DEG pairs. Each SNP-DEG pair was modeled as an eQTL using the interaction of Mtb infection and genotype. Significant genes and eQTLs were defined at FDR < 0.05. Model fit was measured by AIC.

### Data availability

RNA-seq data are available in NCBI dbGaP (phs002445.v1.p1). All scripts and code are available at https://github.com/BIGslu/kimma_MS_public. The kimma software is available at https://github.com/BIGslu/kimma [20].

## Declarations

### Ethics approval and consent to participate

The study protocol was reviewed and approved by the National AIDS Research Committee, The Uganda National Council on Science and Technology, and the institutional review board at University Hospitals Cleveland Medical Center. Written informed consent was obtained from all participants.

### Availability of data and materials

Access to raw transcriptomic data is available through the NCBI database of Genotypes and Phenotypes (dbGaP) accession 002445.v1.p1 (https://www.ncbi.nlm.nih.gov/projects/gapprev/gap/cgi-bin/preview1.cgi?GAP_phs_code=C0TPIcCeWtsG6oqY). Access must be approved by the study’s data access committee (see [8]). R code for analyses in this manuscript are available at https://github.com/BIGslu/kimma_MS_public

## Authors’ contributions

KADM and MCA designed the study. KADM wrote the software, and RMS, BB, TJ, and MSC contributed to software debugging. HMK, WHB, PB, CMS, and TRH provided data. KADM, KM, and SB performed analyses. KADM, KM, and SB wrote the manuscript. All authors reviewed and approved the manuscript prior to submission.

## Acknowledgements

We thank the individual study subjects and their families. We also thank LaShaunda Malone, Keith Chervenak, and Marla Manning for database and program management of the Uganda study as well as the clinical and research staff including Dr. Alphonse Okwera, Dr. Moses Joloba, Hussein Kisingo, Sophie Nalukwago, Dorcas Lamunu, Deborah Nsamba, Annet Kawuma, Saidah Menya, Joan Nassuna, Joy Beseke, Michael Odie, Henry Kawoya, Shannon Pavsek, and Bonnie Thiel

## Supplemental

**Table S1**. Simulated data set analysis sensitivity and specificity at FDR < 0.05. Mean true positive (TP), false positive (FP), false negative (FN), and true negative (TN) detection of differentially expressed genes (DEGs) in simulated data sets of 250 DEG and 750 non-DEG. Sensitivity = TP / (TP + FN) and specificity = TN / (TN + FP).

**Figure S1. Dataset sample designs.**

(A) Simulated differentially expressed gene (DEG) dataset: 100 simulated datasets were created from the media-only condition samples from two clinical groups, RSTR and LTBI. First, 1000 random genes were selected. Then, DEGs were simulated at fold changes from 1 to 100% and 5% error was added. This was repeated with resampling until 100 datasets were obtained. Models were run for both unpaired and paired designs. (B) Mtb vs media real-world data: RNA-seq data was assessed in 49 LTBI individuals to compare Mtb-infected and media conditions in a paired design. (C) RSTR vs LTBI real-world data: RNA-seq data was assessed in Mtb-infected samples to compare 43 RSTR vs 49 LTBI in an unpaired design. For both (B,C), the entire gene dataset with nearly 14,000 protein-coding genes was used. Unrelated and related subsets were defined at kinship < 0.125 (3rd degree) and assessed separately in linear models.

**Figure S2. Runtime to analyze simulated RNA-seq data across unpaired models and software.**

Models were run for 100 simulated datasets of 92 samples and 1000 genes. When available, models were corrected for gene-level weights (kimma, limma) as indicated by panel headers. Multi-processor usage is shown for kimma and DESeq2 at 1 (blue) or 6 processors (orange). Limma does not have this option. Black squares indicate means with standard deviation bars. Across all panels, letters indicate statistically different groups, TukeyHSD FDR < 0.05, in order from shortest time (a) to longest (d).

**Figure S3. Time to run kimma with various models on 6 processors.**

Horizontal red dashed line indicates the base model of ∼condition (Mtb infected vs media) with no additions (none). Sex (female vs male) was used as a covariate and in the interaction term. Contrasts were calculated for all pairwise comparisons of the 4 groups in the condition:sex interaction. The “all preceding” label indicates the model ∼condition + sex + condition:sex + kinship + (1|ptID) with all elements to the left including calculating fit metrics and using gene-level weights. The final “3 models” value summarizes running the “all preceding” model under linear, linear mixed effects, and linear mixed effects with kinship modes, yielding results for each of the 3 models.

**Figure S4. Unpaired design model fit and DEGs in genetically unrelated and related datasets.**

Media samples were subset into unrelated and related subsets at a kinship cutoff of 0.125. Subsets were modeled for RSTR vs LTBI expression. (A) In unrelated individuals, model fit was assessed by AIC with a solid black line indicating a 1:1 fit (e.g. no difference in AIC). Genes better fit without kinship in the model appear above the line (yellow) and those better fit with kinship in the model appear below the line (blue). (B) DEGs were defined at FDR < 0.05. (C,D) Similarly, model fit and DEGs were assessed for the related subset.

