## Supplementary material for "kimma: flexible linear mixed effects modeling with kinship covariance for RNA-seq data": FigureS1

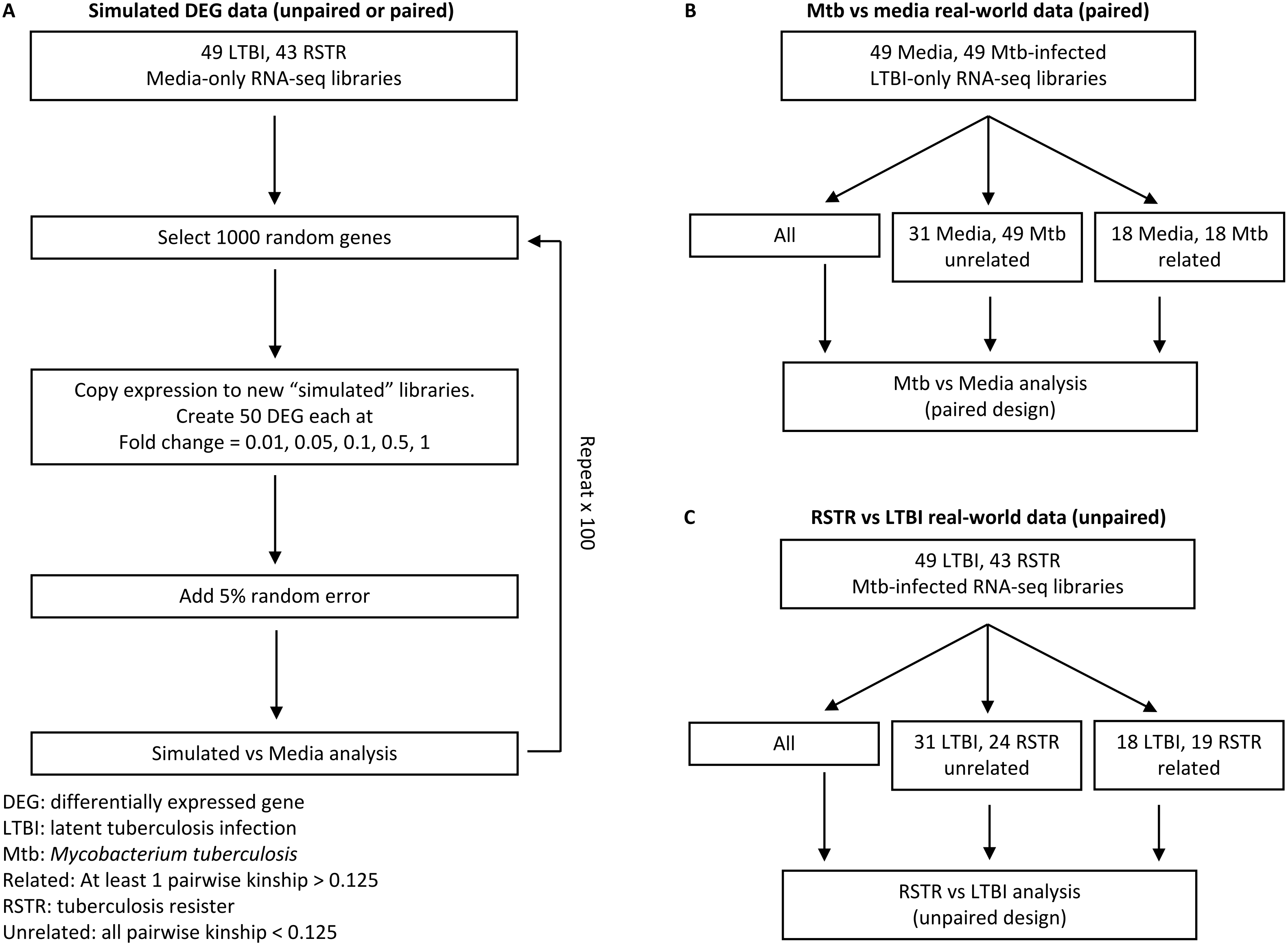


DEG: differentially expressed gene
LTBI: latent tuberculosis infection
Mtb: *Mycobacterium tuberculosis*
Related: At least 1 pairwise kinship > 0.125
RSTR: tuberculosis resister
Unrelated: all pairwise kinship < 0.125
