## Supplementary figures and images for "kimma: flexible linear mixed effects modeling with kinship covariance for RNA-seq data"

### FigureS2

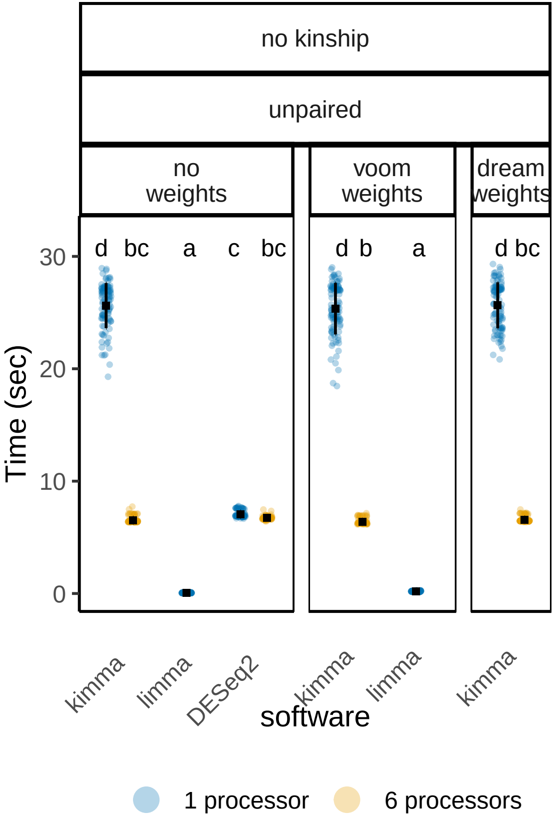

### FigureS3

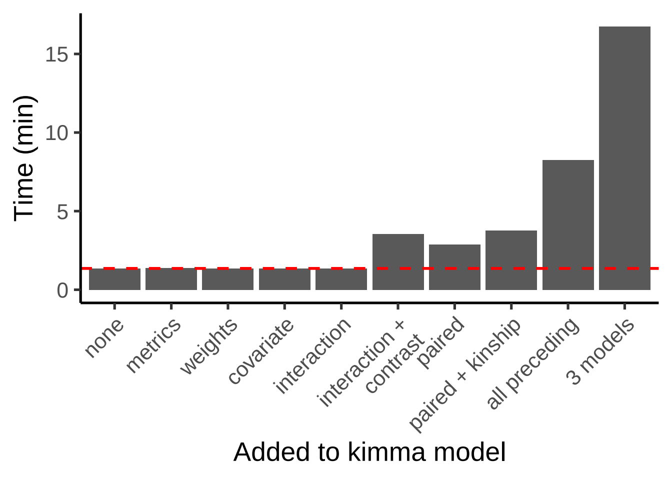

### FigureS4

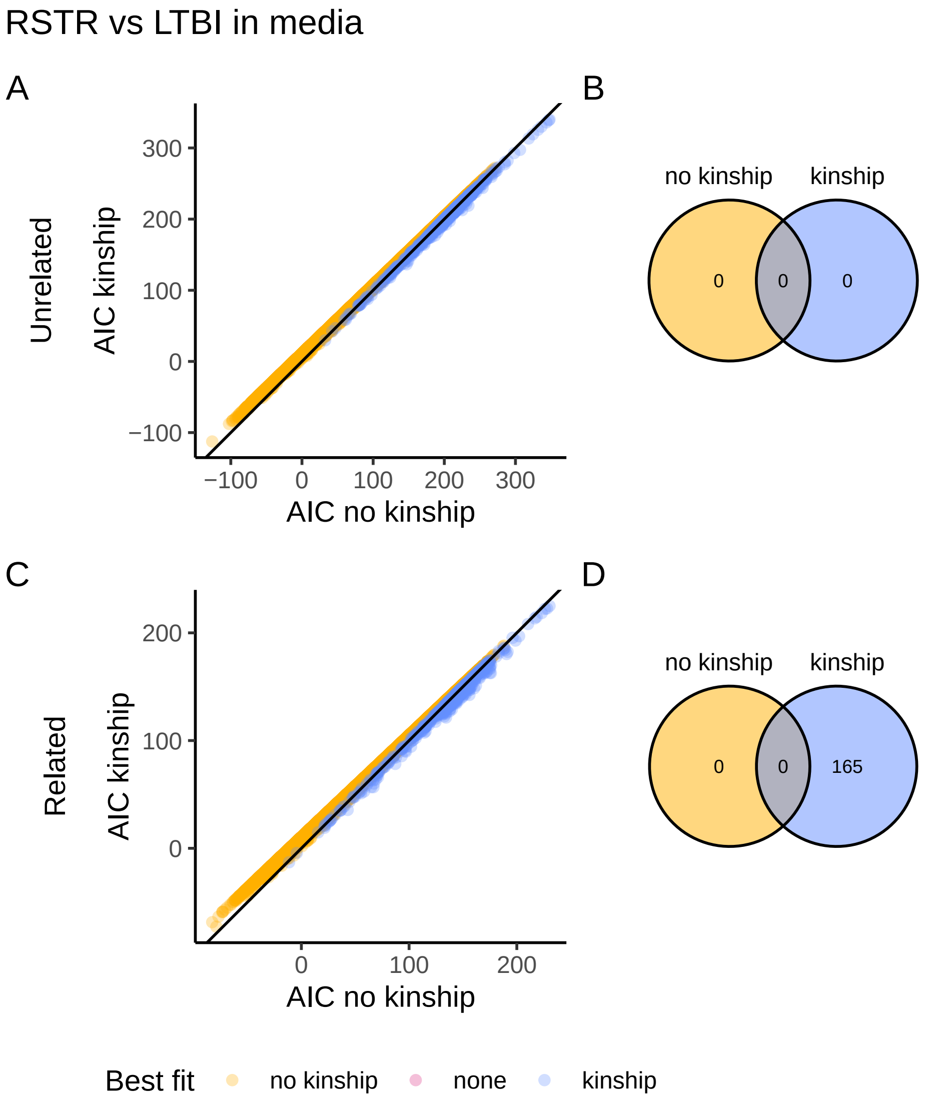
